# Bringing the pancreas patient back to the bench: *Ex vivo* culture of intact human patient derived pancreatic tumour tissue

**DOI:** 10.1101/2020.07.30.223925

**Authors:** John Kokkinos, George Sharbeen, Koroush S. Haghighi, Rosa Mistica C. Ignacio, Chantal Kopecky, Estrella Gonzales-Aloy, Janet Youkhana, Elvis Pandzic, Cyrille Boyer, Thomas P. Davis, Lisa M. Butler, David Goldstein, Joshua A. McCarroll, Phoebe A. Phillips

**Author notes:** **Corresponding author:** Phoebe Phillips, **Affiliation:** Pancreatic Cancer Translational Research Group, School of Medical Sciences, Lowy Cancer Research Centre, UNSW Sydney, NSW, Australia; Australian Centre for Nanomedicine, ARC Centre of Excellence in Convergent Bio-Nano Science and Technology, UNSW Sydney, NSW, Australia. **Email:**.

## Abstract

The poor prognosis of pancreatic ductal adenocarcinoma (PDAC) is attributed to the highly fibrotic stroma and complex multi-cellular microenvironment that is difficult to fully recapitulate in pre-clinical human models. To fast-track translation of therapies and to inform personalised medicine, we aimed to develop a whole-tissue *ex vivo* explant model that maintains viability, 3D multicellular architecture, and microenvironmental cues present in human pancreatic tumours. Patient-derived surgically-resected PDAC tissue was cut into 2 mm explants, cultured on gelatin sponges, and grown for 12 days. Immunohistochemistry revealed that human PDAC tissue explants were viable for 12 days and maintained their original tumour, stromal and extracellular matrix architecture. As proof-of-principle, human PDAC tissue explants responded to Abraxane^®^ treatment with a 3.7-fold increase in cell-death (p=0.0007). PDAC explants were also transfected with polymeric nanoparticles+Cy5-siRNA and we observed abundant cytoplasmic distribution of nanoparticle+Cy5-siRNA throughout the PDAC explant tissue. Our novel model retains the 3D architecture of human pancreatic tumours and has several advantages over standard organoids: presence of functional multi-cellular stroma and fibrosis and no tissue manipulation, digestion, or artificial propagation of organoids. This provides an unprecedented opportunity to study PDAC biology, tumour-stromal interactions and rapidly assess therapeutic response that could drive personalised treatment for PDAC.

## 1. Introduction

Patients with pancreatic ductal adenocarcinoma (PDAC) have less than 9% chance of survival 5 years post-diagnosis ^1^. Despite aggressive treatment regimes, there has been little improvement in patient survival in the past 3 decades ^2,3^. A critical driver of PDAC tumour aggressiveness and a key barrier to drug delivery is the highly fibrotic stroma which can make up the majority of the tumour mass ^4,5^ and is produced by cancer-associated fibroblasts (CAFs) ^6^. A PDAC-CAF cell cross-talk network is known to promote the progression, chemoresistance and metastasis of pancreatic tumours ^6–8^. This is also a key limitation to pre-clinical evaluation of therapeutics for PDAC, as a majority of pre-clinical *in vitro/ex vivo* PDAC models lack the presence of CAFs and an abundant and functional fibrotic stroma. Thus, in order to fast-track the clinical translation of new drug candidates and to identify existing drugs that will be effective on a patient’s individual tumour, there is an unmet need to develop better pre-clinical models that are: 1) easy to establish; 2) cost-effective; 3) provide results in a timely manner to inform patient treatment; 4) avoid mechanical or enzymatic digestion of tissue; and 5) closely reflect the biology of human disease.

Currently, human-PDAC xenograft mouse models and genetically engineered mouse models (GEMMs) are the gold-standard for pre-clinical drug testing and are highly valuable tools to study PDAC biology ^9,10^. Patient-derived xenograft mouse models can reflect inter-patient heterogeneity, but are expensive, time consuming, lack the presence of a functional immune system, and have the extra complexity of infiltrating mouse stroma into a tumour of human origin ^11^. GEMMs contain a complex multicellular fibrotic microenvironment, however the species genome is not identical to the human genome and they are expensive and time-consuming ^9^. Recent years have seen the development of patient-derived organoid models as a pre-clinical model, but these involve the manipulation and mechanical or enzymatic digestion of human tumour tissue and are often derived from a single cell type (tumour cells) ^12,13^ which does not fully mimic human PDAC tumours. Most importantly, organoids often lack a fibrotic stroma and the presence of CAFs and blood vessels. Taken together, these limitations highlight the need to develop more clinically relevant models of PDAC to complement the other models available.

Recent evidence highlights that the *in-situ* spatial interaction between tumour and stromal cells in PDAC can provide clinically-important information ^14^. Thus, an ideal model should reflect the complex microenvironment and contain all cell types within the same 3D architecture as they were present in a pancreatic tumour. Here, we have developed a new PDAC pre-clinical model that retains the 3D architecture of human patient derived PDAC tumours. Importantly, this model does not involve any chemical, enzymatic, or mechanical digestion of PDAC tissue and thus avoids artificially skewing cell populations. We further demonstrate that this model can be applied to test both clinically approved chemotherapy drugs as well as novel therapeutics including a nano-based gene silencing drug developed in our lab. This new model provides a unique opportunity to closely study the biology of the PDAC tumour microenvironment, to identify novel gene targets and test new treatment strategies in a cost-effective and timely manner.

## 2. Results

### 2.1. Culture and characterisation of human patient derived PDAC explants

Human PDAC tumour tissue was obtained from patients undergoing surgical resection (pancreaticoduodenectomy) of pancreatic cancer. Patient characteristics are listed in **Table 1**. A small piece of tissue was resected from the tumour mass and transported within 15 minutes to the laboratory on ice. The tumour tissue was cut into 2 mm explants and placed on pre-soaked gelatin sponges in triplicate (**Figure 1a** and **Supplementary Fig. S1 online**). A control explant was fixed immediately as the day 0 timepoint and cultured explants were fixed on days 5, 7, 9 and 12 post-establishment of the model. H&E staining revealed that both tumour and stromal architecture of the explants from 3 different PDAC patients was retained throughout the 12-day culture (**Figure 1b** and **Supplementary Fig. S2 online**). With similar architecture to the uncultured day 0 controls, we observed in patient 1 explants an arrangement of both cytokeratin-positive tumour cells and stromal α-SMA-positive CAFs that were Ki67 positive and TUNEL negative, thus confirming their viability throughout the 12-day culture (**Figure 2a**). Picrosirius red staining demonstrated that the explants retained an abundant distribution of fibrillar collagen throughout the 12 days of culture (**Figure 2a**). Similarly, patient 2 and 3 explants showed consistent cytokeratin, α-SMA and Ki67 staining across all the timepoints from days 0-12 (**Supplementary Fig. S3-S4 online**). We also showed that TUNEL and collagen staining is comparable for patient 2 and 3 explants at days 0 and 12 (**Supplementary Fig. S5 online**).

**Table 1:** Patient and tumour characteristics. Staging based on American Joint Committee on Cancer 8^th^ Edition. Metastasis category not assessable by histology.

| Patient ID | Age (yr) | Sex | Tumour type | Histological grade | Stage |
| --- | --- | --- | --- | --- | --- |
| Patient 1 | 77 | Female | Pancreatic ductal adenocarcinoma | 3 | T2N2 |
| Patient 2 | 86 | Female | Pancreatic ductal adenocarcinoma | 2 | T2N2 |
| Patient 3 | 67 | Male | Pancreatic ductal adenocarcinoma with loss of MSH6 expression | 2 | T3N2 |
| Patient 4 | 50 | Male | Pancreatic neuroendocrine tumour | 1 | T2N0 |
| Patient 5 | 56 | Female | Pancreatic neuroendocrine tumour | 2 | T3N1 |
| Patient 6 | 50 | Female | Metastatic leiomyosarcoma to the pancreas | N/A | N/A |

**Figure 1.**
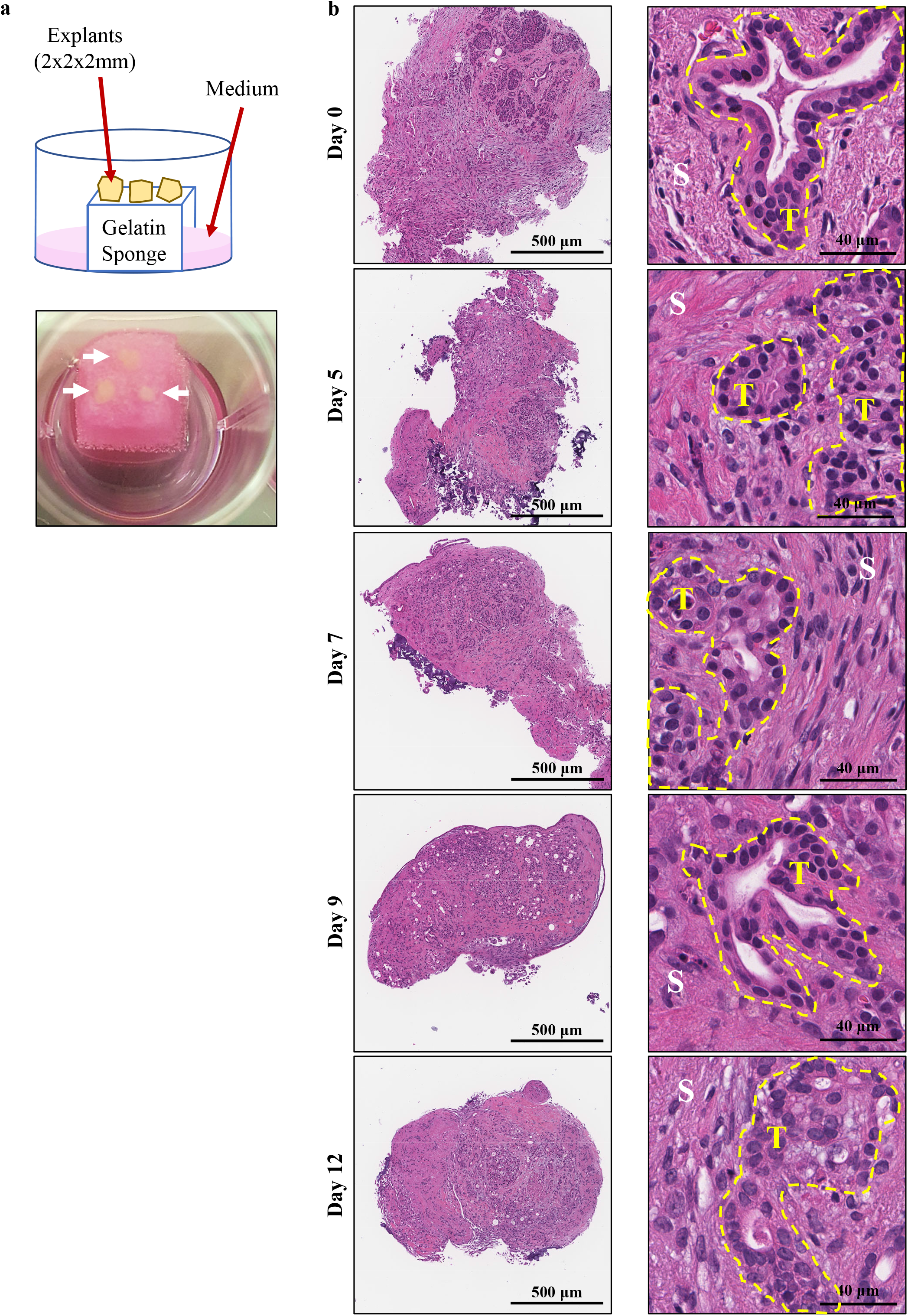
Tumour and stromal architecture are maintained in human patient derived PDAC explants for 12 days of culture. **(a)** Schematic showing the set-up of the *ex vivo* culture method and a representative photo of a gelatin sponge containing 3 PDAC explants in a well of a 24-well plate. White arrows point to the 3 explants on the gelatin sponge during culture. **(b)** Representative H&E images of patient 1 explants at low and high magnification from days 0-12. Tumour elements outlined in yellow and compartments labelled as tumour (T) and stroma (S).

**Figure 2.**
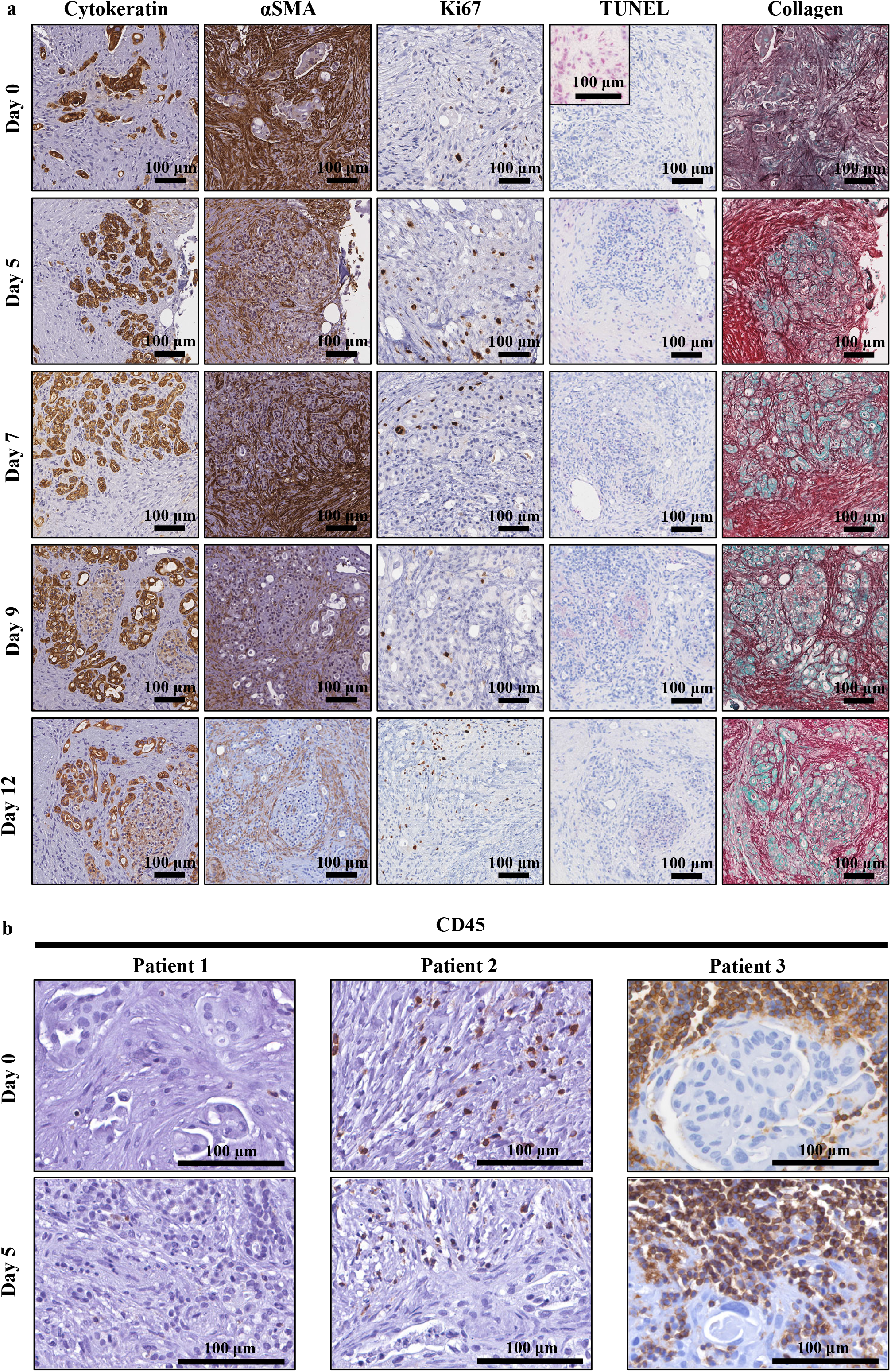
Characterisation of human patient derived PDAC explants cultured for 12 days. **(a)** Immunohistochemistry was performed for cytokeratin, α-smooth muscle actin (αSMA), Ki67, TUNEL and collagen (picrosirius red/methyl green) on patient 1 explants from days 0-12. Small insert in TUNEL staining shows positive control (DNAse treated). **(b)** CD45 immunohistochemistry in PDAC explants from patients 1, 2 and 3 at day 0 and day 5 of culture.

We also performed immunohistochemistry for CD45 (marker for leukocytes) and observed a high degree of lymphocyte infiltrate in the day 0 control explants of patient 3, and this was maintained in explants cultured for 5 days (**Figure 2b**). Interestingly, according to the pathology report of the surgically resected tumour, patient 3 had a rare loss of MSH6 expression which could explain the unusually high amount of CD45-positive immune cells in the tumour. In contrast, patient 2 explants had much fewer CD45-positive lymphocytes while patient 1 explants had almost no CD45-positive cells. Notably, the amount of immune infiltrate across all three patients was maintained between day 0 and day 5 explants. Explants cultured for 7-12 days had very few CD45-positive cells (data not shown), suggesting that the lymphocytes present in the patient’s tumours only remain viable for 5 days in our *ex vivo* explant model.

We also performed light sheet microscopy, demonstrating the ability to image a whole tissue explant in 3D. The 3D reconstruction of the day 0 control tissue explant showed individual cell nuclei with distinct areas of F-actin (phalloidin) expression throughout the explant (**Supplementary Fig. S6 online**). This could suggest a dispersed arrangement of F-actin rich stromal cells ^15^ throughout the tissue and provides an opportunity to stain for other cell markers and image their distribution in 3D.

In addition to patients with PDAC, we also cultured explants from 2 patients with pancreatic neuroendocrine tumours as well as a patient with a rare metastasis of a leiomyosarcoma to the pancreas. H&E staining demonstrated that the architecture of these explants was also maintained after 12 days of culture (**Supplementary Fig. S7 and S8 online**). Overall, these findings show that the *ex vivo* explant model can maintain the viability, cell composition and extracellular matrix of tissue explants from a range of human pancreatic tumours including PDAC and pancreatic neuroendocrine tumours for at least 12 days of culture.

### 2.2. Testing of clinical and novel therapeutics in pancreatic tumour tissue explants

We next investigated whether the *ex vivo* explant culture model can be used to test clinically approved or novel therapeutics. We first tested Abraxane^®^ (human albumin-bound paclitaxel) which is currently one of the chemotherapeutic agents used in first-line therapy for PDAC ^16^. Abraxane^®^ was added to the medium reservoir every 3 days and allowed to penetrate through the gelatin sponge into the tissue explant via capillary action. We performed TUNEL staining to assess levels of cell death. Abraxane^®^ treated tissue explants had a significantly higher percentage of TUNEL positive cells compared to untreated control explants (3.7-fold increase, p=0.0007, n=3; **Figure 3a-b**). H&E staining revealed architectural changes to tumour cells with Abraxane^®^ treatment including fragmentation of ductal elements (black arrows, **Figure 3a**).

**Figure 3.**
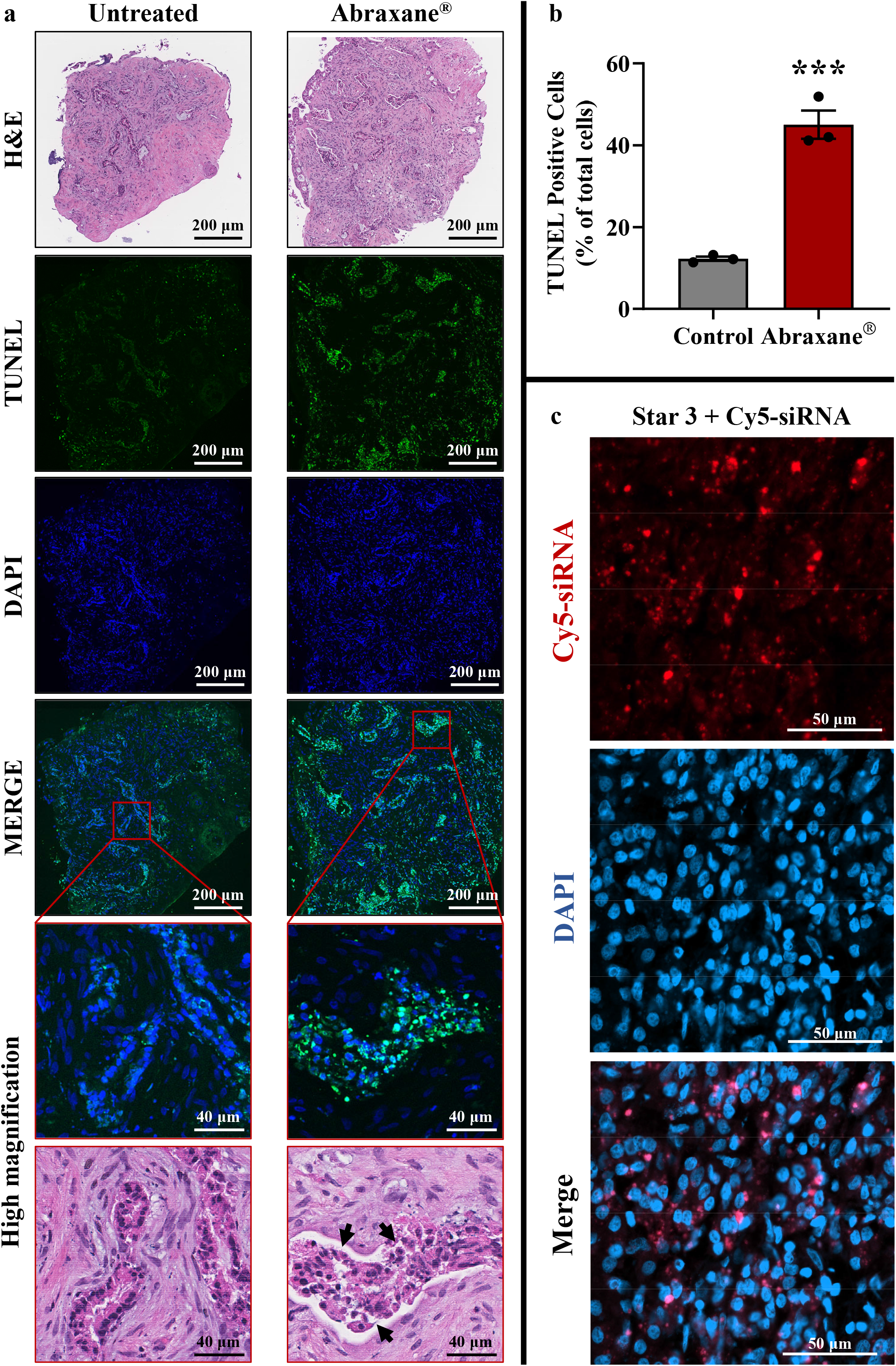
Testing of clinical and novel therapeutics in human patient derived PDAC explants. **(a)** Patient-derived explants were treated with or without 150ng/mL Abraxane^®^ on days 0, 3, 6 and 9, then fixed on day 12. Representative H&E and TUNEL staining. Black arrows point to areas of ductal fragmentation in Abraxane^®^ treated explants. **(b)** Quantification of TUNEL positive cells using QuPath demonstrated increased cell death in Abraxane^®^ treated explants compared to untreated controls. An unpaired t-test was performed to compare TUNEL-positivity in control vs Abraxane^®^ treated explants from a single patient, with 3 explants per treatment from distinct regions of the patient’s tumour. Bars represent mean ± S.E.M., ***p=0.0007. **(c)** Representative images showing uptake of Cy5-siRNA coupled to polymeric nanoparticles (Star 3) in human pancreatic tumour PDAC explants after 24 hours of treatment.

Another key potential of our novel *ex vivo* explant model is the ability to perform real-time mechanistic studies and study cell-cell interactions in an unmanipulated piece of human PDAC tissue. We hypothesised that gene therapeutics such as siRNA would be a useful tool to perform such mechanism studies, hence we examined whether our tissue explant model is amenable to transfection with nanoparticle-siRNA ^17,18^. A di-block co-polymer nanoparticle (Star 3) developed in our lab was complexed to fluorescently labelled siRNA. Human PDAC explants were obtained from a patient and placed in triplicate on gelatin sponges that had been pre-soaked in medium containing Star 3 complexed to siRNA. After 24 hours, tissue explants were processed for frozen tissue sections and Star 3-siRNA uptake assessed by confocal microscopy. As proof-of-principle, we demonstrated that Star 3-siRNA treated PDAC tissue explants had abundant uptake of fluorescent siRNA throughout the PDAC tissue explants (**Figure 3c**), thus providing rationale to test siRNA knockdown of therapeutic targets in future studies.

## 3. Discussion

In this study, we describe the development and characterisation of a novel human pre-clinical model of pancreatic ductal adenocarcinoma (PDAC) that maintains the viability, 3D multicellular architecture and microenvironmental cues of unmanipulated patient derived tumours. This simple and cost-effective model provides unprecedented opportunity to closely study the biology of pancreatic cancer, to identify novel gene therapeutic targets for tumour and stromal cells, to test the anti-cancer activity of novel drugs or combination treatments, and has potential to inform precision medicine for pancreatic cancer. PDAC urgently requires new and more effective treatments. However, for the efficacy of new therapeutic strategies to be evaluated, there is a need for robust pre-clinical models that accurately reflect the biology of the disease in patients. Currently, a broad range of PDAC models are available; ranging from cell lines, GEMMs, patient derived xenografts, and most recently, organoid cultures ^9,10,13^. There is no perfect model and each model has its strengths and limitations. For example, 2D *in vitro* cell line cultures can provide important insights into the signalling pathways dysregulated in cancer cells, but they lack the complexity of a 3D tumour mass. Subcutaneous or orthotopic mouse models of PDAC have the advantage of a 3D arrangement of cancer cells but are often derived from a single cell type. Patient derived xenografts can model interpatient heterogeneity, but are time consuming, expensive, have the complexity of infiltrating mouse stromal cells into a tumour of human origin, and the poor tumour engraftment can potentially bias more aggressive disease ^11^. GEMMs have the advantage of spontaneously forming tumours that can mimic patient tumourgenicity in an immune proficient setting but are also time consuming, expensive, and may not always reflect therapeutic response due to mouse and human species differences ^9^. Recently, patient-derived organoids have gained much attention for their ability to rapidly model interpatient heterogeneity, but they often consist of tumour cells that have been dissociated from their native extracellular matrix and tumour microenvironment ^12,13^. Additionally, most organoids lack the characteristic fibrotic stroma, which is a key drug delivery barrier and is known to drive the progression of PDAC. With this in mind, we identified that there is a significant gap in the pre-clinical models available for PDAC, and in particular, there is a need for a model with: 1) abundant and functional stroma, and 2) multicellular architecture with the same 3D organisation as present in human disease.

At the onset of this study, we hypothesised that the *ex vivo* human patient-derived explant model, established in prostate and breast cancer, has the potential to overcome many of these limitations ^19^. This model involves the culture of freshly resected tumour tissue explants (around 1-2 mm diameter) on a gelatin sponge support scaffold soaked in culture media ^19^.

Nutrients can be taken up by the explants via capillary action through the gelatin sponge, preventing the need to submerge the tissue in culture media which often leads to its degradation within a few days. This model has been used to culture prostate, breast and ovarian cancer explants, but has yet to be translated to PDAC ^19–23^.

Recently, a study by Misra et al (2019) ^24^ cultured 350 μm thick PDAC tumour tissue-slices on a cell-culture Millicell^®^ insert for up to 96 hours. While this was the first study to culture whole-tissue slices of PDAC and represents significant progress in developing more clinically relevant models of PDAC, the culture was only maintained for 96 hours and increasing levels of tissue death were observed from as early as 24 hours of culture ^24^. A follow up study demonstrated that these cultured PDAC tissue slices have genomic stability with minimal transcriptome changes throughout the 72 hours of culture ^25^. Several other studies have developed *in vitro* co-culture models containing tumour cells, cancer-associated fibroblasts (CAFs), and endothelial cells ^26–30^, but these models do not perfectly mimic the *in situ* complexity and heterogeneity of human PDAC tumours. For example, an impressive study by Gupta et al ^26^ developed a coculture model of cancer cells, endothelial cells and CAFs on a polyurethane scaffold. While such models can be useful for high-throughput screening of therapeutics, they consist of cells artificially dissociated from their native architecture and microenvironment which may not represent the complexity of human disease.

Here, we report for the first time that the gelatin sponge *ex vivo* explant culture method can be used to culture human patient derived PDAC whole-tissue tumour explants for up to 12 days. We cultured human PDAC explants derived from patient tumours immediately following surgical resection of the tumour. Remarkably, both the tumour and stromal architecture of the explants were retained throughout the 12-day culture, and this was reproduced across 3 PDAC patients. Indeed, the tissue explants showed dispersed cytokeratin-positive tumour elements surrounded by a dense arrangement of αSMA-positive CAFs, and this organisation was highly comparable from days 0-12. Picrosirius red staining showed an abundant network of fibrillar collagen throughout the explants. While PDAC is generally characterised by an immunosuppressive microenvironment with limited intratumoural immune cell populations, we cultured explants from a patient with a rare loss of MSH6 expression (patient 3) where we observed a prominent infiltrate of CD45-positive lymphocytes. Loss of MSH6 results in a deficiency in mismatch repair which can increase neoantigen presentation on tumour cells and promote T-cell infiltration ^31^. Notably, this immune population was maintained up to 5 days of culture, thus providing a window to potentially study the effects of immunotherapy on a 3D culture of human PDAC tissue. There is also potential to add patient autologous immune cells to the explant culture and study how they interact with different cell types and the effects of stromal remodelling on immune cell activity and invasion.

Interestingly, we demonstrated that our model can be used to culture pancreatic neuroendocrine tumour explants as well as tissue explants derived from a patient with metastatic leiomyosarcoma to the pancreas, providing an opportunity to study other pancreatic malignancies which also have poor patient outcome.

The ability to maintain the viability of a patient’s tumour in the lab over a 12-day window is a major advancement and has the potential to revolutionise pancreatic cancer research. Firstly, this model provides an unprecedented opportunity to perform real-time mechanistic studies to better understand interactions of tumour cells with their surrounding stromal and immune cell populations in the same native 3D architecture as they were present in a patient’s tumour. We also propose that the *ex vivo* tissue explant model can become an integral component of the drug development pipeline as it allows for the potential of new drugs to be evaluated within the context of the major cell types present in an individual patient’s tumour. Inter-patient heterogeneity can be recapitulated in the explant model, allowing the applicability of a new drug to be assessed and to inform which patients would likely benefit from a proposed new treatment. Analysis of secreted factors in the medium reservoir also provides an opportunity to study potential circulating biomarkers as a measure of treatment response.

With this in mind, we investigated whether our tissue explant model is amenable to transfection with nano-based gene silencing drugs. We showed that polymeric nanoparticles which can deliver siRNA to PDAC cells both *in vitro* and *in vivo*, delivered siRNA throughout the human PDAC tumour explants. No obvious toxicity from the nanoparticle treatment was observed. Thus, the potential to administer therapeutics to patient-derived explants in this model may provide an opportunity to evaluate a novel therapeutic agent before using expensive and time-consuming mouse models. From a nanomedicine perspective, this model holds great potential to test the interaction of nanoparticles with all cell types present in a tumour, and to observe their biodistribution in a clinically relevant 3D piece of tumour tissue. This information is critical, especially in PDAC, to facilitate the translation of nanomedicines to the clinic ^32^. In addition, the potential to transfect patient derived human PDAC tumour explants with siRNA or other gene modifying (e.g. miRNA mimics / inhibitors, DNA or CRISPR/Cas) agents provides an opportunity to perform real-time mechanistic studies in tumour tissue containing a 3D multicellular architecture and complex microenvironment.

Most importantly, the PDAC explant model has potential to inform a personalised medicine program for PDAC. Patient-derived xenograft models are being investigated as a tool to guide personalised medicine in other cancer types ^33^, but the length of time required to establish these models makes it unsuitable for cancers such as PDAC that have such poor survival. Recent years have seen the focus shift to organoid cultures to inform individual patient treatment ^12,34–38^, but a limitation of most organoid models is that they lack the presence of stroma – a major drug delivery barrier for PDAC. While organoids may reflect tumour cell intrinsic resistance to a given chemotherapy agent, a patient’s tumour cells deemed to be “sensitive” to a chemotherapeutic may in fact be resistant in the presence of a fibrotic stroma containing CAFs and immune cells that are well known to cross-talk with tumour cells to promote chemoresistance.

As a proof-of-principle, we demonstrated that patient-derived tissue explants can be treated with Abraxane^®^ and anti-cancer activity measured by TUNEL staining as a readout for cell-death. We will next test a broader range of chemotherapeutics and prospectively assess whether response in our explant model correlates with patient response in the clinic. If successful, this will pave the way for our explant model to drive a personalised medicine program for PDAC. Patient tumour explant culture can be established following surgical resection, and explants cultured in the presence of a range of chemotherapy drugs to determine a sensitivity profile to these chemotherapeutic agents. Within a short timeframe (approximately 3-4 weeks including analysis), this information can then be fed back to the patient’s clinician to inform personalised treatment.

Importantly, this model is both cost effective and time efficient. There is no need to add complex and expensive growth factors to the culture medium and the gelatin sponge scaffold used in our model is commonly used by dentists and is thus readily available. Preparation of the explants can be completed in approximately 30 minutes after receiving the surgical sample, so a team of two researchers could process explants from multiple patients within a given week. In our experience, we were able to obtain at least 30 explants from each patient which could allow around 10 different treatment groups to be established from each patient with 3 explants per treatment group to ensure tumour heterogeneity is reflected.

In addition to informing precision medicine, our PDAC tumour explant model may also provide insight into how chemotherapy treatment affects the microenvironment of PDAC tumours. This has been studied using the same model in human prostate cancer explants and revealed patient-specific changes to the stroma and microenvironment following chemotherapy treatment ^39^. A recent study demonstrated that adjuvant chemotherapy increased the mutational burden of patient PDAC tumours compared to treatment naïve tumours ^40^. This raises the question whether chemotherapy treated PDAC tumours with a higher mutational burden have increased neoantigen expression and would potentially benefit from immunotherapy ^41^. Such questions can be addressed by our explant model with the possibility of testing the effects of chemotherapy on the microenvironment of patient whole-tissue PDAC explants.

Another potential of the PDAC explant model is the opportunity to study the effects of stromal reprogramming strategies for PDAC treatment. The concept of normalising the stroma has recently become a popular strategy to improve drug delivery in PDAC. For example, vitamin D receptor inhibition has been shown to induce CAF quiescence and improve gemcitabine efficacy in mouse tumours ^42^. Our PDAC explant model provides a unique opportunity to assess the therapeutic potential of such stromal reprogramming agents by evaluating the effects on CAFs and the extracellular matrix in a model that uniquely maintains the tumour and stromal architecture of patient derived tissue.

While the pancreatic tumour explant model established in this study holds promise for PDAC research, a limitation is that we have only collected tumour samples from patients with surgically resectable disease which represents approximately 15-20% of all PDAC patients and these patients have a better prognosis compared to those with unresectable PDAC ^2,43^. Future studies should focus on developing and strengthening collaborations between scientists and clinicians with the aim to obtain fresh tumour tissue from unresectable patients through biopsy of their primary tumour and/or metastatic tumours.

Nonetheless, this study has established a promising new pre-clinical model of PDAC that retains the native 3D multicellular architecture of human pancreatic tumours over 12 days of culture. While we do not believe that this model replaces the need for mouse models, we propose that the *ex vivo* tissue explant model is a clinically relevant complement to currently available pre-clinical models. Importantly, the *ex vivo* tissue explant culture method can answer fundamental biological questions and has potential to guide a precision medicine program for PDAC.

## 4. Material and Methods

### 4.1. Ex Vivo Explant Culture

Prior to collection of tumour tissue, haemostatic gelatin sponges (Johnson & Johnson, Cat. JJ-12505) were briefly submerged in culture medium containing high-glucose DMEM, 10% FBS, 5 mM GlutaMAX, 0.01 mg/mL hydrocortisone, 0.01 mg/mL insulin and 1x antibiotic/antimycotic solution (all from Sigma-Aldrich). Once sponges were soaked, they were placed in 24-well plates and 500 μL culture medium was added to each well to cover the bottom-half of each sponge. The 24-well plate was placed in a 37°C/5% CO_2_ incubator until explants were ready for culture.

De-identified tumour samples were obtained from patients undergoing a pancreaticoduodenectomy (Whipple procedure) at Prince of Wales Hospital or Prince of Wales Private Hospital, Randwick, NSW, Australia. All patients provided informed consent through the Health Science Alliance Biobank, all work was approved by UNSW human ethics (HC180973) and all experiments were performed in accordance with the relevant guidelines and regulations. Tumour tissue was transported to a tissue culture hood on ice within 15 minutes of receiving the sample. The tissue was immediately placed in ice-cold PBS containing 1x antibiotic/antimycotic solution. First, any regions of normal pancreas or fat tissue were dissected away from the tumour sample. The remaining tumour tissue, which could be identified by its more solid texture, was cut into 3 pieces of equal size. The 3 pieces were placed in separate petri dishes labelled as “L”, “M”, or “R”. Each of these 3 pieces were then cut into 2×2×2 mm explants using a scalpel. If tumour tissue was limited, smaller explants were prepared at 1 mm^3^. Explants from the “L” piece were all placed on the bottom left corner of each pre-soaked gelatin sponge, explants from the “M” piece placed in the middle region, and explants from the “R” piece placed in the bottom right corner as described in **Supplementary Fig. S1 online**. This ensured that each gelatin sponge contained explants from 3 distinct regions of the tumour tissue to account for intratumoural heterogeneity. A single explant from each of the “L”, “M”, and “R” pieces was immediately fixed in 4% paraformaldehyde and was designated the day 0 control. Once all explants had been placed on the sponges, the 24-well plate was then placed in a 37°C 5% CO_2_ incubator. On average, approximately 30-50 explants could be obtained from each patient, although this varied depending on how much tumour tissue was obtained from surgery. Culture medium was replaced daily, with fresh insulin, hydrocortisone and antibiotic/antimycotic solution added to the medium immediately prior to it being added to the culture plate. Explants were fixed in 4% paraformaldehyde at the indicated time points. Fixed explants were embedded in paraffin and 5 μm slices were prepared at the Histopathology Service at the Garvan Institute of Medical Research. All explants were stained with Haematoxylin and Eosin (H&E) for visualisation of explant architecture.

Patient-derived explants were treated with or without Abraxane^®^ (Specialised Therapeutics Australia; 150 ng/mL) by adding Abraxane^®^ to the medium used to pre-soak the gelatin sponges on day 0 and 500 μL medium containing Abraxane was added to each well. Abraxane treatment was repeated on days 3, 6 and 9, and medium replaced with fresh medium on all other days. The 150 ng/mL dose of Abraxane^®^ was calculated to match a dose of 10 mg/kg typically used in mouse models, assuming an average 20 g mouse has a blood volume of 1.2 mL.

### 4.2. Immunohistochemistry of patient-derived pancreatic tumour explants

Patient-derived explant tissue sections were stained for cytokeratin (DAKO, Cat. M3515; 1:100 overnight at 4°C), α-smooth muscle actin (αSMA) (Sigma, Cat. A5228; 1:1000 1-hour incubation at room temperature) or CD45 (ThermoFisher Scientific, Cat. 14-9457-82; 1:100 overnight at 4°C). Briefly, tissue sections were deparaffinised at 60°C for 30 minutes then rehydrated through consecutive washes in xylene, ethanol, and water. Antigen retrieval was performed by microwaving slides for 4 minutes in 10mM citrate buffer + 0.05% Tween-20 at pH6.0, followed by a 30-minute incubation at 104°C. Non-specific peroxidase activity was blocked with 1% hydrogen peroxide + 1% methanol for 10 minutes at room temperature. Non-specific antibody binding was blocked with 10% goat serum in PBS for 1 hour at room temperature. After blocking, tissue samples were stained with primary antibodies as indicated above and isotype control antibodies were used as negative controls. Biotinylated anti-rabbit secondary antibody (Vector laboratories, Cat. BA-1000) was used at 1:100 for 45 minutes at room temperature followed by a 5-minute incubation with Vectastain^®^ ABC kit (Vector laboratories). 3,3’ diaminobenxidine (DAB) was used as the substrate, and tissues were counterstained with hematoxylin. Ki67 staining (ThermoFisher Scientific, Cat. RM-9106; 1:1000 dilution) was performed on a LeicaBond Autostainer. All stained tissue sections were scanned on an AperioXT (Leica Biosystems) or Vectra Polaris (PerkinElmer) slide scanner.

TUNEL staining was performed according to manufacturer’s instructions (Sigma, Cat. 11684809910). Tissue sections were deparaffinised at 60°C for 30 minutes then rehydrated through consecutive washes in xylene, ethanol, and water. Antigen retrieval was performed by with 20 μg/mL proteinase K for 15 minutes at 37°C. Positive control slides were treated with 3 U/mL DNase-1 (New England Biolabs, Cat. M0303) for 10 minutes at room temperature. Tissue sections were incubated for 1 hour at 37°C with TUNEL enzyme solution and label solution diluted 1:2 in TUNEL dilution buffer. For assessment of TUNEL fluorescence (as performed for Abraxane^®^ treatment experiment), tissue sections were mounted with ProLong™ Gold antifade mounting medium with DAPI (Invitrogen, Cat. P36931) and fluorescence scanned on a Vectra Polaris (PerkinElmer) slide scanner. QuPath software was used to quantify the amount of TUNEL-positivity as a percentage of DAPI-positive cells. An unpaired t-test (GraphPad Prism 8) was performed to compare TUNEL-positivity in control vs Abraxane^®^ treated explants from a single patient, with 3 explants per treatment from distinct regions of the patient’s tumour. For the remainder of tissue sections, TUNEL staining was followed by a 30-minute incubation with alkaline phosphatase converter for 30 minutes at 37°C. Fast Red (abcam, Cat. ab64254) was used as the substrate, and tissue sections were counterstained with haematoxylin.

### 4.3. Picrosirius red staining of collagen

Patient-derived explant tissue sections were stained with 0.1% picrosirius red for fibrillar collagen and counterstained with methyl green through the UNSW Mark Wainwright Analytical Centre Biomedical Imaging Facility (UNSW Sydney).

### 4.4. Uptake of polymeric nanoparticle-siRNA complexes in human patient derived PDAC tumour explants

Polymeric nanoparticles (Star 3) were synthesised as previously described ^18^. Star 3 nanoparticles (62.5 μg) were complexed for 5 minutes with 25 μg Cy5-labelled siRNA (Dharmacon custom Luc2 siRNA; Cy5-GCUUAGGCUGAAUACAAAUUUU). The complexed Star 3 + Cy5-siRNA was added to the medium used to soak the gelatin sponges immediately prior to addition of the explants to the sponges and 500 μL of medium containing Star 3 + Cy5-siRNA was then added to each well. After 24 hours, explants were embedded and frozen in Tissue-Tek^®^ Optimal Cutting Temperature Compound (OCT; VWR International). OCT-embedded sections (10 μm thick) were mounted with ProLong™ Gold antifade mounting medium with DAPI (Invitrogen, Cat. P36931). Fluorescent siRNA uptake was visualised on a Zeiss 800 confocal microscope by taking at least 3 representative images from each explant.

### 4.5. Light-sheet microscopy

Fixed human PDAC explants were first embedded in agarose to provide the support and simplify the mounting of the sample for light-sheet microscopy. Delipidation and optical clearing of the explants was performed as previously described ^44^. Briefly, the tissues were incubated in reagent 1 [25wt% urea, 25wt% N,N,N’,N’-tetrakis (2-hydroxypropyl) ethylenediamine and 15wt% Triton X-100] for 7 days at 37 °C. Next, the explants were washed in PBS for 3x 1 hour to remove the excess reagent. Optically cleared samples were then stained with Alexa Fluor 647-conjugated F-actin antibody (1:200, ThermoFisher, USA) and DAPI (1:200,ThermoFisher, USA) in PBS for 3 days at 37 °C, followed by 3x 1 hour washes in PBS. The samples were then equilibrated in reagent 2 [50wt% sucrose, 25wt% urea, 10wt% 2,2,2′-nitrilotriethanol and 0.1% (v/v) Triton X-100] for 2 days. The stained explants were imaged on a light-sheet microscope (Zeiss Light-sheet Z.1; Carl Zeiss; Germany) operated at the Biomedical Imaging Facility (Mark Wainwright Analytical Centre, UNSW Sydney). The imaging was performed with a 5 × / 0.16 detection lens and two light-sheet illuminations (left and right) using 5 × / 0.1 illumination lenses. For each illumination (left and right) a separate image was captured for each channel imaged. The fluorescent signal was collected using a 460–500 nm emission filter and 405 nm excitation laser for DAPI and 660/LP nm emission filter and 638 nm excitation laser for Alexa Fluor 647 F-actin. To capture the entire explant, a Z stack of ∼1000 frames at 30 ms exposure on two CMOS (PCOEdge) cameras with 1920 × 1920 pixels images (2.329 μm pixel size and 4 μm optical sectioning steps) was used. Zen software (Carl Zeiss; Germany) was used to combine the left and right illumination images at each Z-plane. Further data processing, rendering of Z-stacks and visualisation in 3D was performed using Imaris software (Andor Technology; Switzerland).

## 6. Acknowledgements

Biospecimens and data used in this research were obtained from the HSA Biobank, UNSW Biorepository, UNSW Sydney, Australia. We sincerely thank the patients who kindly consented to donate their tumour samples for research, without whom this would not have been possible. We would like to thank Dr Carmel Quinn and Dr Anusha Hettiaratchi of the HSA Biobank for their support in managing the clinical samples and patient consent. We also acknowledge the Biomedical Imaging Facility within the Mark Wainwright Analytical Centre (UNSW Sydney) and the Histopathology Service at the Garvan Institute of Medical Research. We would also like to acknowledge our collaborating community consumers Mr Gino Iori and Ms Claire Harvey for their invaluable input.

## Funding

This research was made possible by a Tour de Cure PhD Support Scholarship (Kokkinos, Phillips, Goldstein, RSP-011-18/19) and a Cancer Institute NSW ‘The Professor Rob Sutherland AO Make a Difference Award’ (Goldstein, 2017/AWD002). The following sources supported author contributions and research: Australian Government Research Training Program Scholarship and UNSW Sydney Scientia PhD Scholarship (Kokkinos), NHMRC project grant (Phillips, McCarroll, Goldstein, APP1144108) and the Avner Pancreatic Cancer Foundation Innovation Grant (Phillips, McCarroll, Goldstein, Davis and Sharbeen, APCF0050618), Cancer-Institute NSW ECF/CDFs (Sharbeen, CDF181166), Olivia Lambert Foundation (McCarroll), Cancer Australia (Phillips, McCarroll and Goldstein, APP1126736) and Beat Cancer Project Principal Cancer Research Fellowship (Butler, PRF1117).

## 7. Author Contributions

JK, GS, DG, JM and PP wrote the manuscript. JK, GS, KH, RI, CK, EG, JY, and EP performed the experiments. All authors contributed to the design and interpretation of experiments and reviewed the manuscript.

## 8. Additional Information

The authors declare no conflict of interest. All data has been made available in the results section of the text, the figures, and supplementary figures.

## Notes

### Competing Interest Statement

The authors have declared no competing interest.

