## Supplementary Figures for "Bringing the pancreas patient back to the bench: *Ex vivo* culture of intact human patient derived pancreatic tumour tissue"

*[1]Pancreatic Cancer Translational Research Group, School of Medical Sciences, Lowy Cancer Research Centre, UNSW Sydney, NSW, Australia; [2]Australian Centre for Nanomedicine, ARC Centre of Excellence in Convergent Bio-Nano Science and Technology, UNSW Sydney, NSW, Australia; [3]Prince of Wales Hospital, Prince of Wales Clinical School, UNSW Sydney, NSW, Australia; [4]Biomedical Imaging Facility, Mark Wainwright Analytical Centre, Lowy Cancer Research Centre, UNSW Sydney, NSW, Australia; [5]Australian Centre for Nanomedicine, UNSW Sydney, NSW, Australia; [6]Centre for Advanced Macromolecular Design, School of Chemical Engineering, UNSW Sydney, NSW, Australia; [7]ARC Centre of Excellence in Convergent Bio-Nano Science and Technology and Australian Institute for Bioengineering and Nanotechnology, The University of Queensland, Queensland, Australia;, [8]ARC Centre of Excellence in Convergent Bio-Nano Science and Technology, Monash Institute of Pharmaceutical Sciences, Monash University, VIC, Australia; [9]Adelaide Medical School and Freemasons Foundation Centre for Men's Health, University of Adelaide, SA, Australia; [10]South Australian Health and Medical Research Institute, Adelaide, SA, Australia; [11]Children's Cancer Institute, Lowy Cancer Research Centre, UNSW Sydney, NSW, Australia; [12]School of Women's and Children's Health, UNSW Sydney, NSW, Australia.*
***Corresponding author**: Phoebe Phillips

### Supplementary Figures

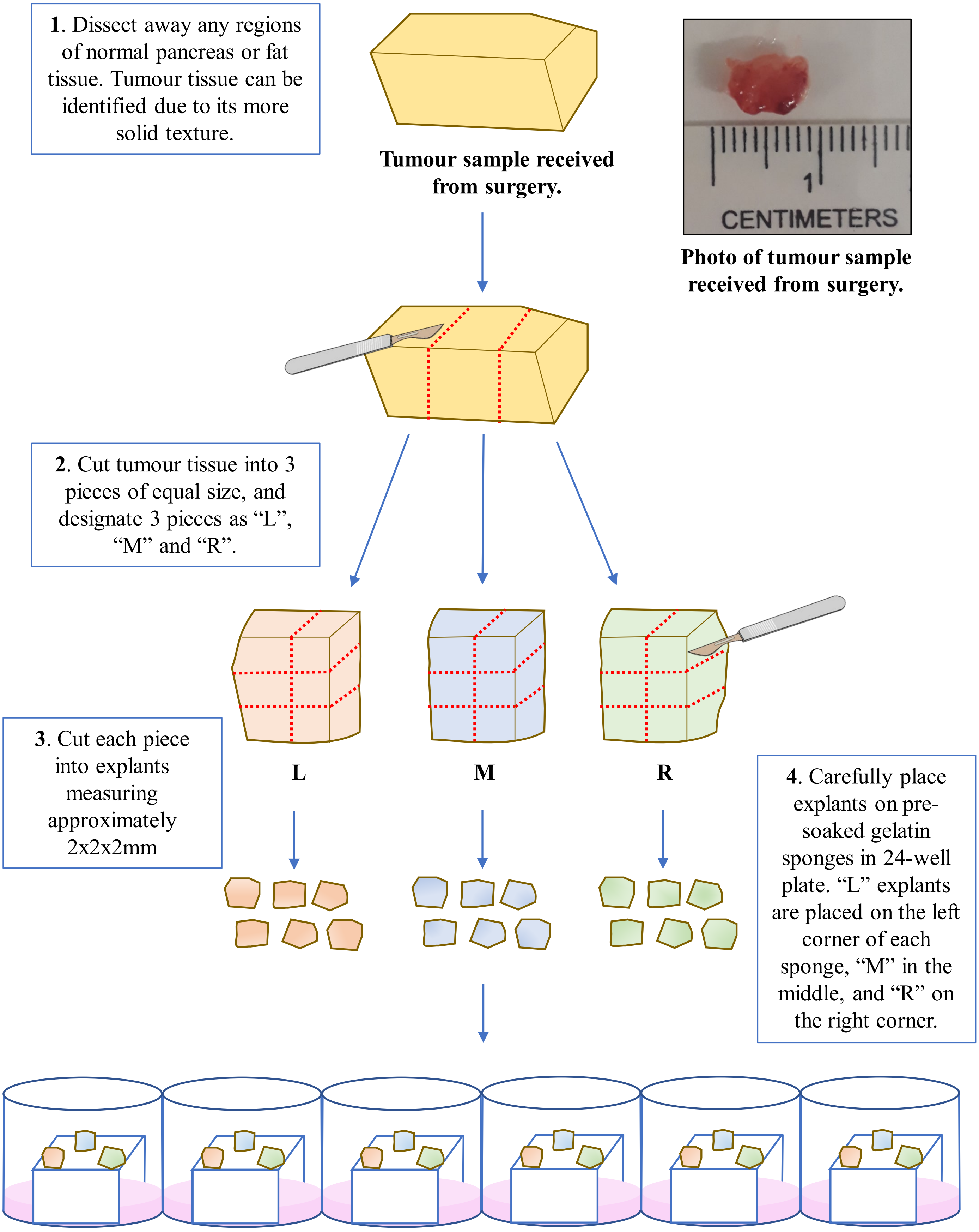


**Supplementary Figure S1. Preparation of patient derived PDAC tumour explants from surgically resected tissue.** Workflow diagram showing how PDAC tumour tissue is prepared and cut for explant culture.

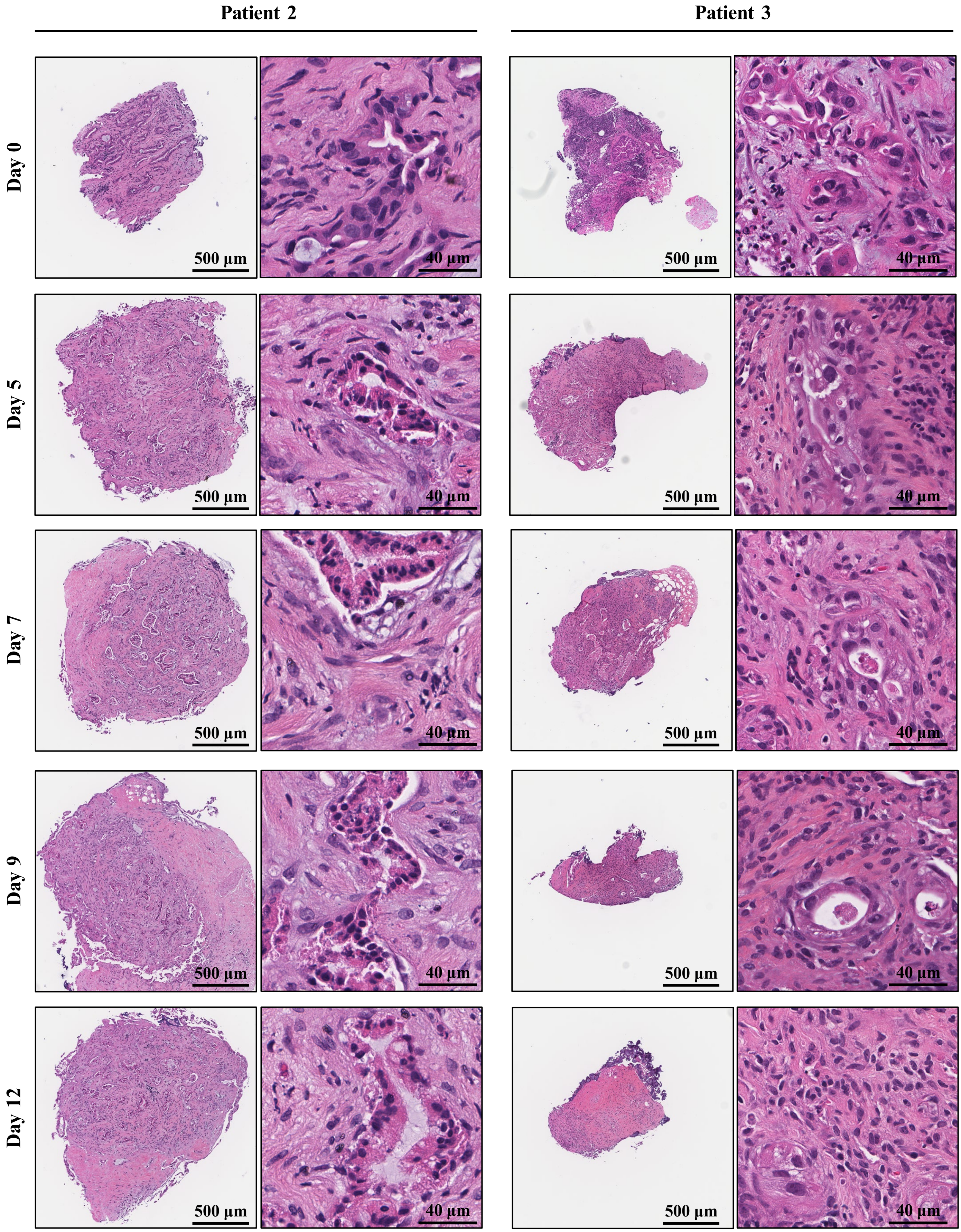


**Supplementary Figure S2. H&E staining of human PDAC explants from patients 2 and 3.** Representative H&E images of patient 2 and 3 explants at low and high magnification from days 0-12.

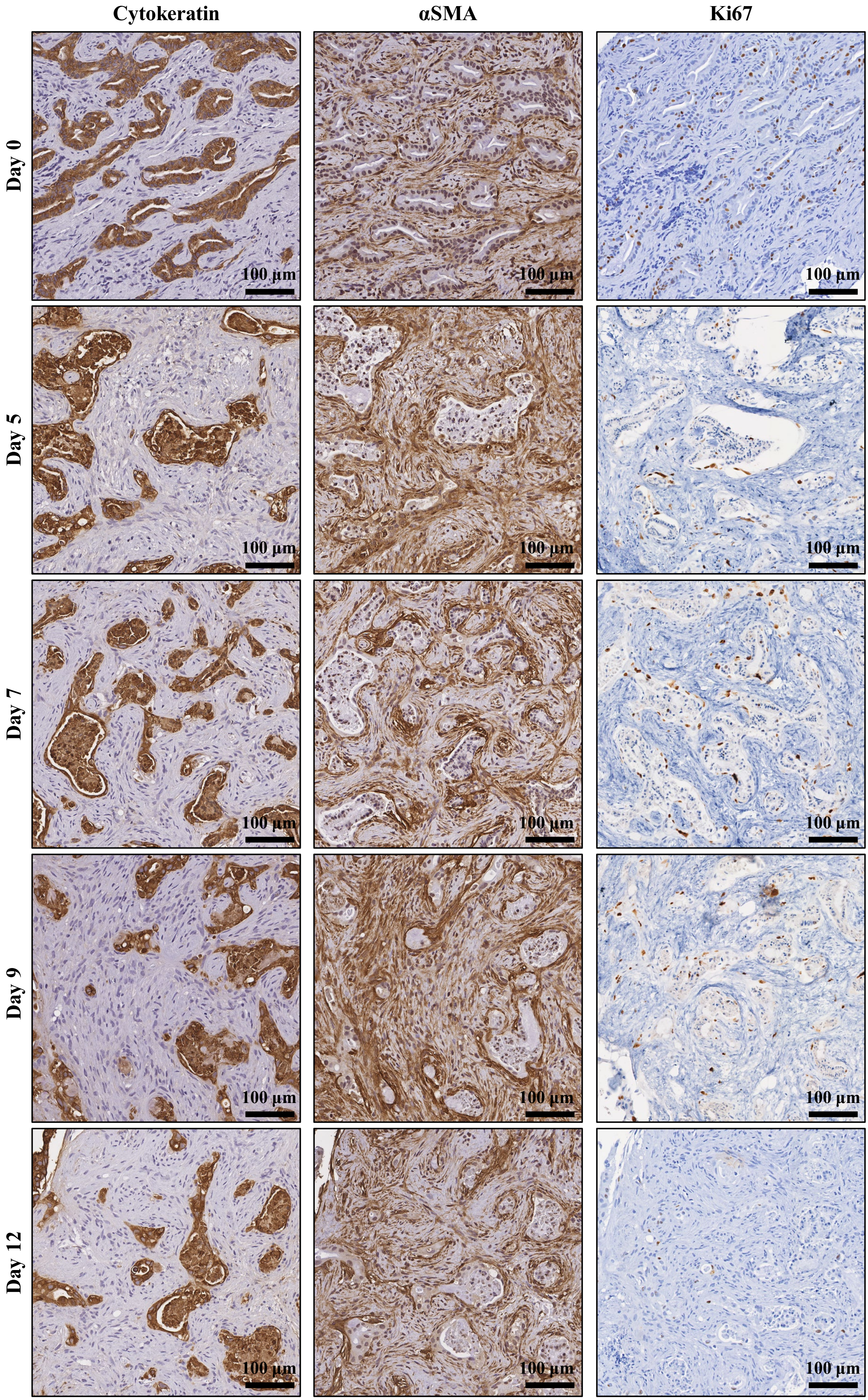


**Supplementary Figure S3. Characterisation of patient 2 human PDAC explants from days 0-12.**

**Supplementary Figure S3. Characterisation of patient 2 human PDAC explants from days 0-12.** Immunohistochemistry was performed for cytokeratin, α-smooth muscle actin (αSMA) and Ki67 on patient 2 PDAC explants from days 0-12.

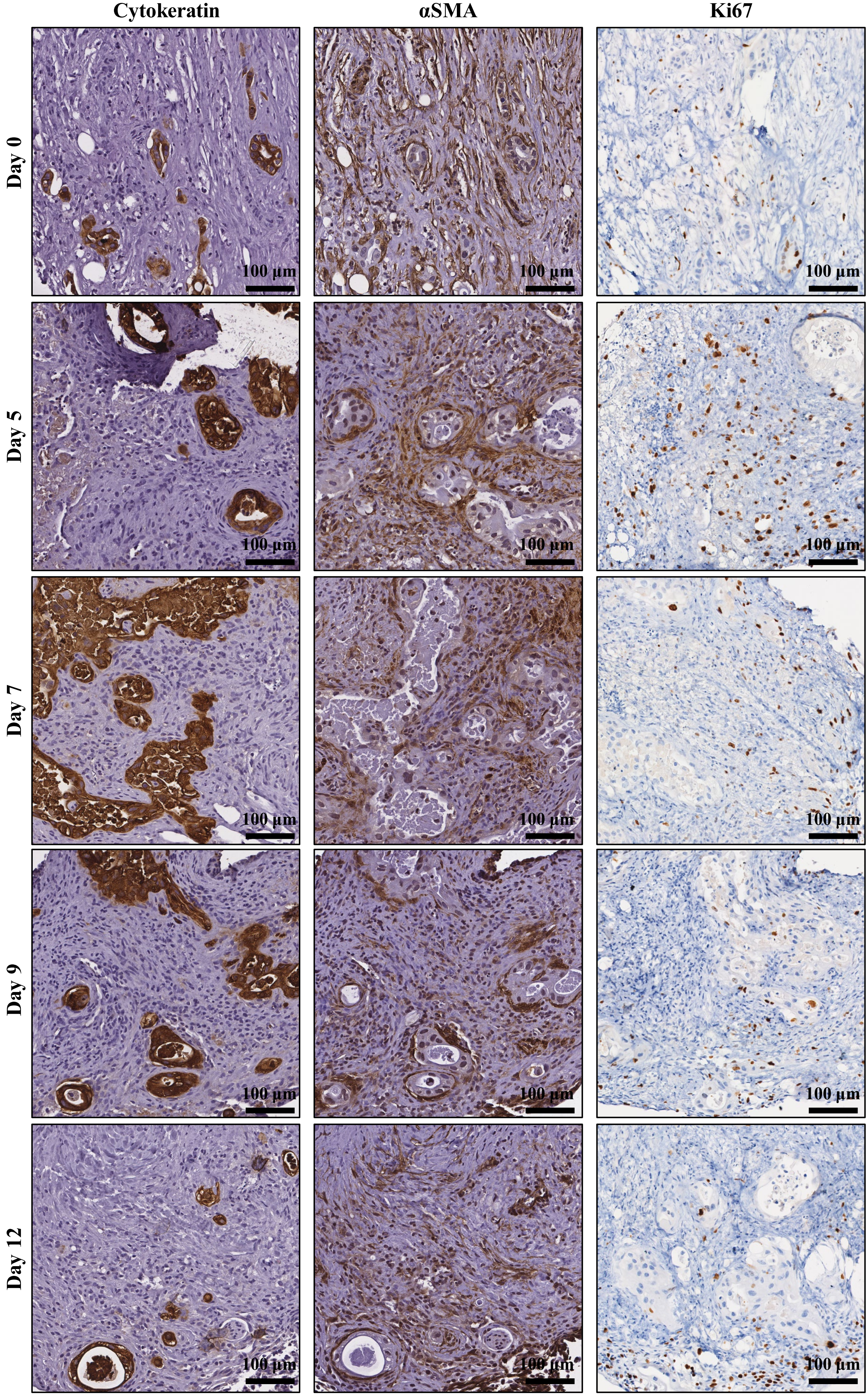


**Supplementary Figure S4. Characterisation of patient 3 human PDAC explants from days 0-12.**

**Supplementary Figure S4. Characterisation of patient 3 human PDAC explants from days 0-12.** Immunohistochemistry was performed for cytokeratin, α-smooth muscle actin (αSMA) and Ki67 on patient 3 PDAC explants from days 0-12.

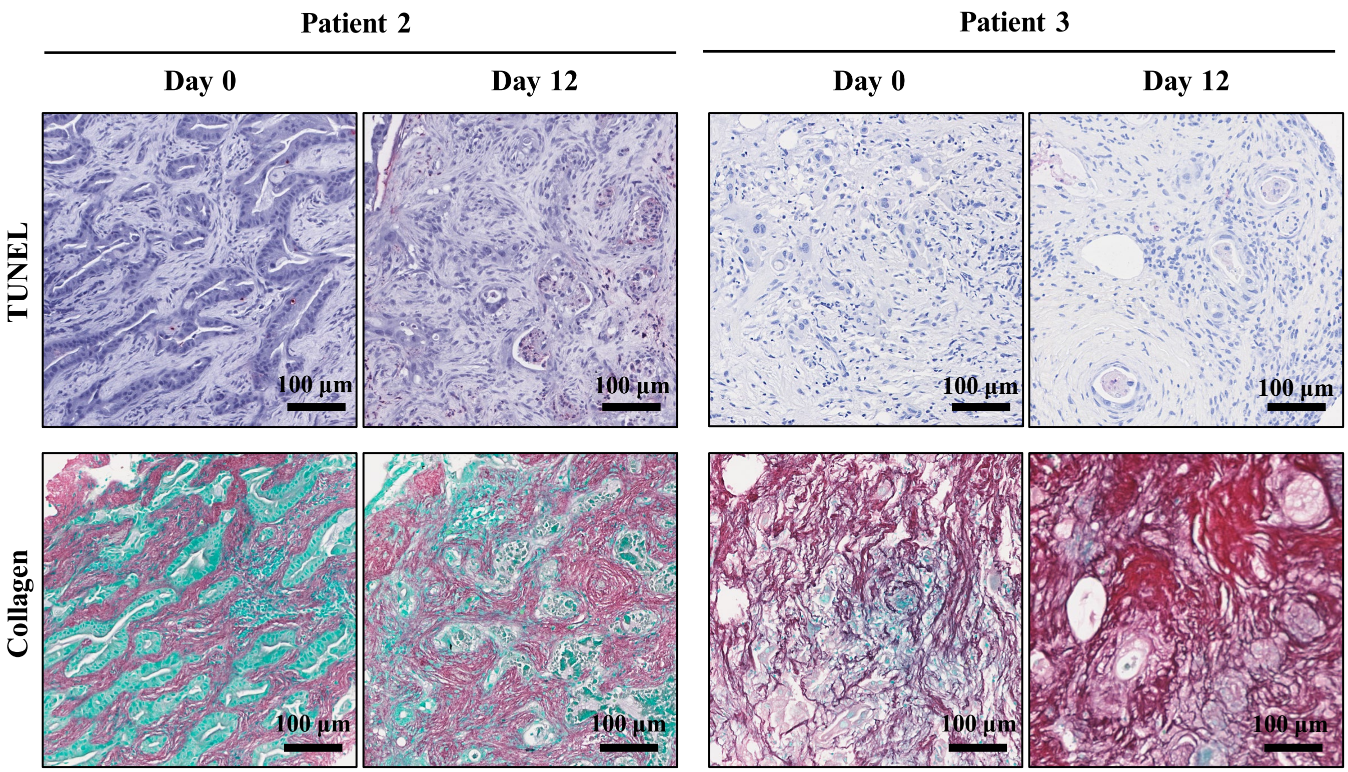


**Supplementary Figure S5. TUNEL and collagen staining of patient 2 and 3 human PDAC explants at day 0 and 12.** TUNEL and collagen (picrosirius red/methyl green) staining was performed on patient 2 and 3 PDAC explants at days 0 and 12. All scale bars represent 100 µm.

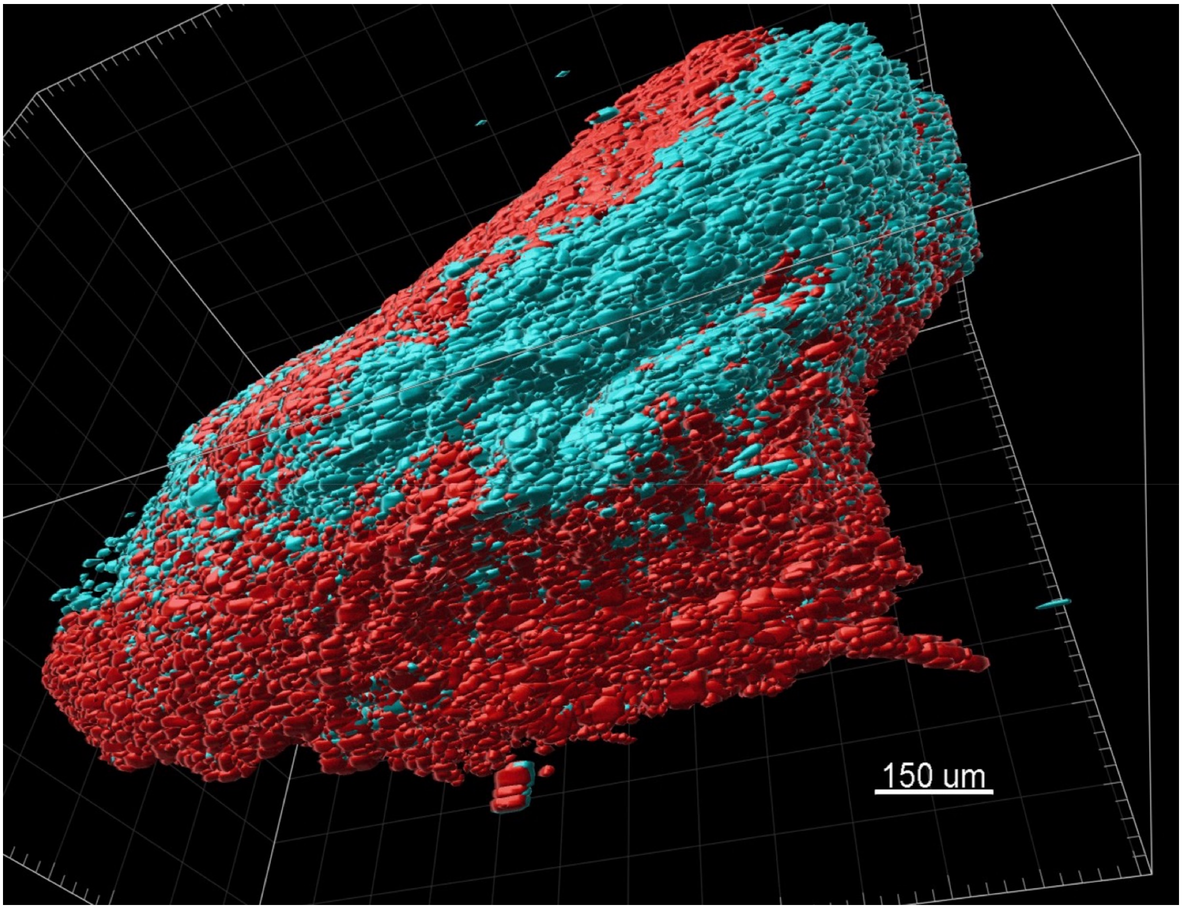


**Supplementary Figure S6. 3D light-sheet microscopy imaging of a human PDAC explant**. 3D reconstruction of the whole-tissue explant showing single cell nuclei (cyan) and F-actin (red) rich stromal cells.

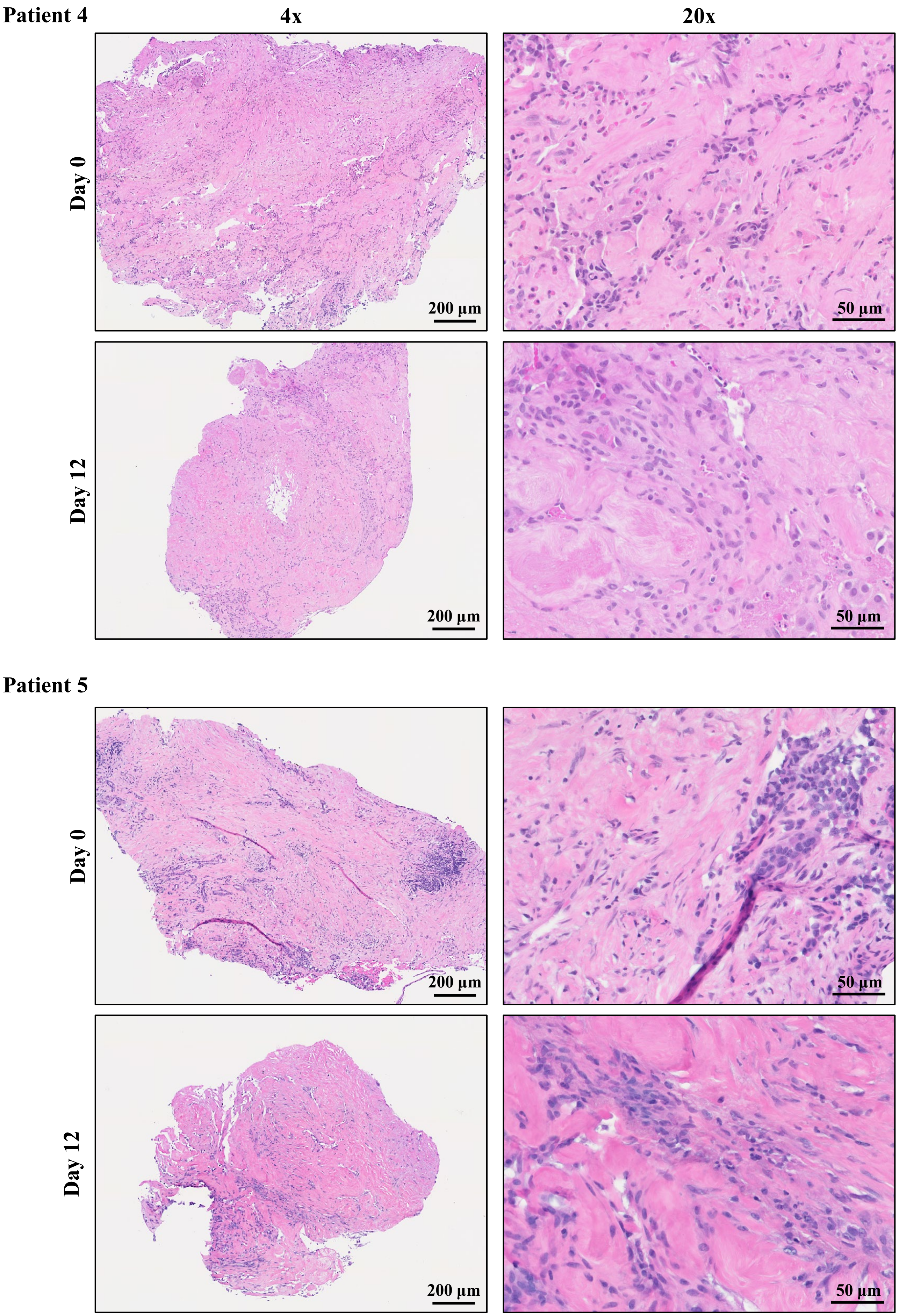


**Supplementary Figure S7. Human explant culture of pancreatic neuroendocrine tumours from 2 patients.** Representative images of H&E staining of pancreatic neuroendocrine tumours at day 0 and day 12 of culture.

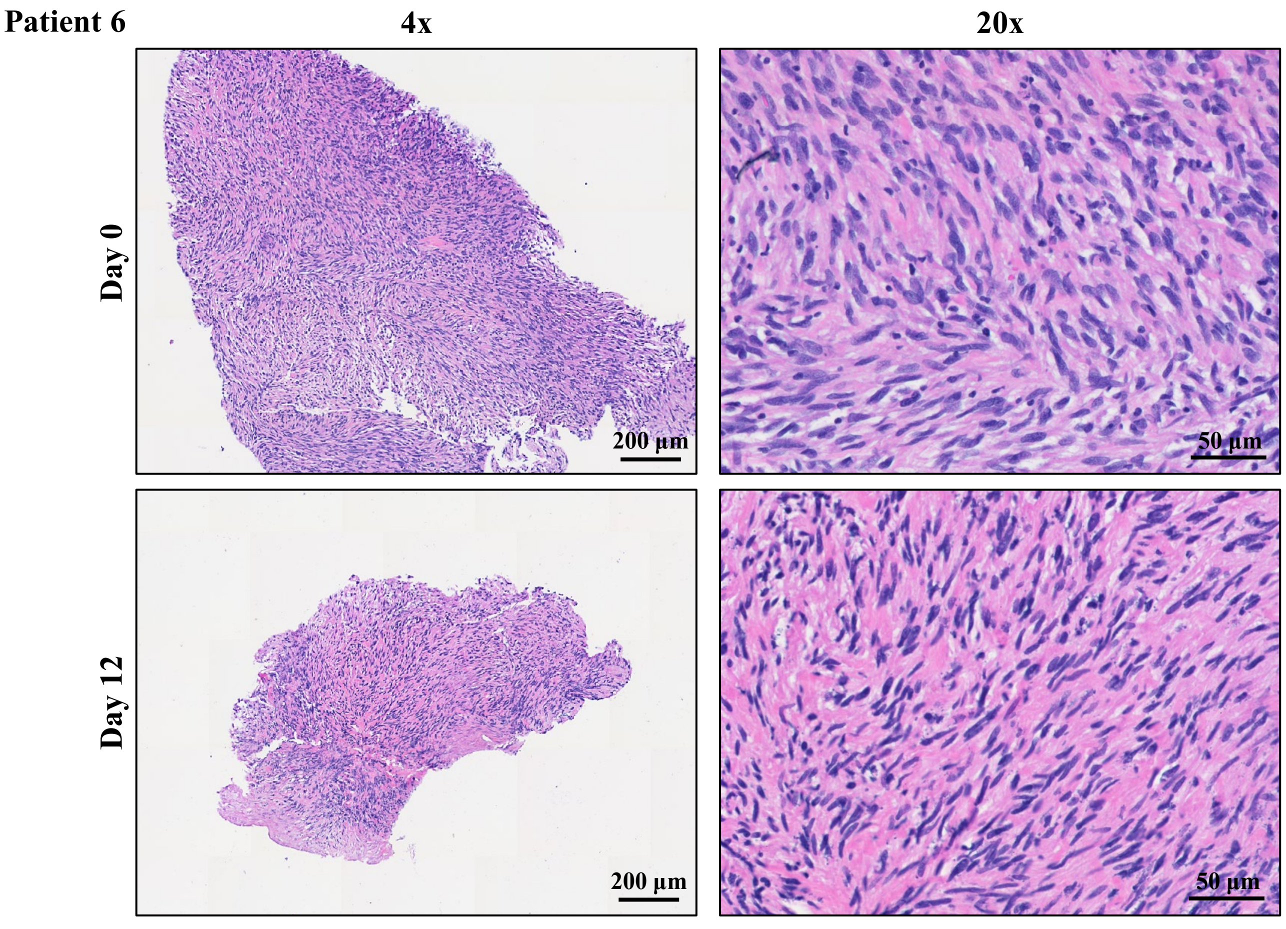


**Supplementary Figure S8. Human explant culture of a rare metastatic leiomyosarcoma metastasis to the pancreas.** Low and high magnification representative images of H&E staining of a metastatic leiomyosarcoma to the pancreas at day 0 and day 12 of culture.
